# Pangolin homology associated with 2019-nCoV

**DOI:** 10.1101/2020.02.19.950253

**Authors:** Tao Zhang, Qunfu Wu, Zhigang Zhang

## Abstract

To explore potential intermediate host of a novel coronavirus is vital to rapidly control continuous COVID-19 spread. We found genomic and evolutionary evidences of the occurrence of 2019-nCoV-like coronavirus (named as Pangolin-CoV) from dead Malayan Pangolins. Pangolin-CoV is 91.02% and 90.55% identical at the whole genome level to 2019-nCoV and BatCoV RaTG13, respectively. Pangolin-CoV is the lowest common ancestor of 2019-nCoV and RaTG13. The S1 protein of Pangolin-CoV is much more closely related to 2019-nCoV than RaTG13. Five key amino-acid residues involved in the interaction with human ACE2 are completely consistent between Pangolin-CoV and 2019-nCoV but four amino-acid mutations occur in RaTG13. It indicates Pangolin-CoV has similar pathogenic potential to 2019-nCoV, and would be helpful to trace the origin and probable intermediate host of 2019-nCoV.

---

In the late December of 2019, an epidemic pneumonia (the W.H.O. announced it recently as Corona Virus Disease (COVID-19)(*1*)) outbreak in city of Wuhan in China and soon widely spreads all over the world. According to authoritative statistics, the COVID-19 has caused more than 40,000 laboratory-confirmed infections with more than 1000 deaths by 12 February 2020 and is still increasing. It was caused by a novel identified coronavirus 2019-nCoV (the International Committee on Taxonomy of Viruses (ICTV) renamed this virus as severe acute respiratory syndrome coronavirus 2, SARS-CoV-2 (*1*)). Released complete genomes of 2019-nCoVs (*2, 3*) have helped rapid identification and diagnosis of the COVID-19. Another key task is to find potential origin of 2019-nCoV(*4*). Unsurprisingly, like SARS-CoV and MERS-CoV(*5*), the bat is still a probable origin of the 2019-nCoV because the 2019-nCoV shared 96% whole genome identity with a bat coronavirus Bat-CoV-RaTG13 from *Rhinolophus affinis* from Yunnan Province(*2*). However, SARS-CoV and MERS-CoV usually pass into medium host like civets or camels before leaping to humans(*4*). It indicates that the 2019n-Cov was probably transmitted to humans by some other animals. Considering the earliest COVID-19 patient reported no exposure at the seafood market(*6*), finding intermediate host of 2019-nCoV is vital to block its transmission.

On 24 October 2019, Liu and his colleagues from the Guangdong Wildlife Rescue Center of China (*7*) firstly detected the existence of SARS-liked coronavirus from lung samples of two dead Malayan Pangolins with a frothy liquid in lung and pulmonary fibrosis, which is close on the outbreak of COVID-19. From their published results, all virus contigs assembled from 2 lung samples (lung07, lung08) showed not high identities ranging from 80.24% to 88.93% with known SARS coronavirus. Hence, we conjectured that dead Malayan pangolin may carry a new coronavirus close to 2019-nCoV.

To confirm our assumption, we downloaded raw RNA-seq data (SRA accession number PRJNA573298) of those two lung samples from SRA and conducted consistent quality control and contamination removing as described by Liu’s study(*7*). We found 1882 clean reads from lung08 sample mapped upon 2019-nCoV reference genome (GenBank Accession MN908947)(*3*) with high genome coverage of 76.02%. We performed *de novo* assembly of those reads and totally obtained 36 contigs with length ranging from 287bp to 2187bp with mean length of 700bp. Blasting against proteins from 2845 coronavirus reference genomes including RaTG13, 2019-nCoVs and other known coronaviruses, we found 22 contigs can be best matched to 2019-nCoVs (70.6%-100% aa identity; average: 95.41%) and 12 contigs matched to Bat SARS-like coronavirus (92.7%-100% aa identity; average: 97.48%) (Table S1). These results indicate Malayan pangolin indeed carries a novel coronavirus (here named as Pangolin-CoV) close to 2019-nCoV.

Using reference-guided scaffolding approach, we created Pangolin-CoV draft genome (19,587 bp) based on the above 34 contigs. Remapping 1882 reads against the draft genome resulted in 99.99% genome coverage at a mean depth of 7.71 X (range: 1X-47X) (**Figure 1A**). Based on Simplot analysis, Pangolin-Cov showed highly overall genome sequence identity throughout the genomes to RaTG13 (90.55%) and 2019-nCoV (91.02%) (**Figure 1B**), although there is greatly high identity 96.2% between 2019-nCoV and RaTG13(*3*). Another SARS-like coronavirus more similar to Pangolin-Cov were Bat SARSr-CoV ZXC21(85.65%) and Bat SARSr-CoV ZC45 (85.01%). These results indicate Pangolin-Cov may be the common origin of 2019-nCoV and RaTG13.

**Figure 1.**
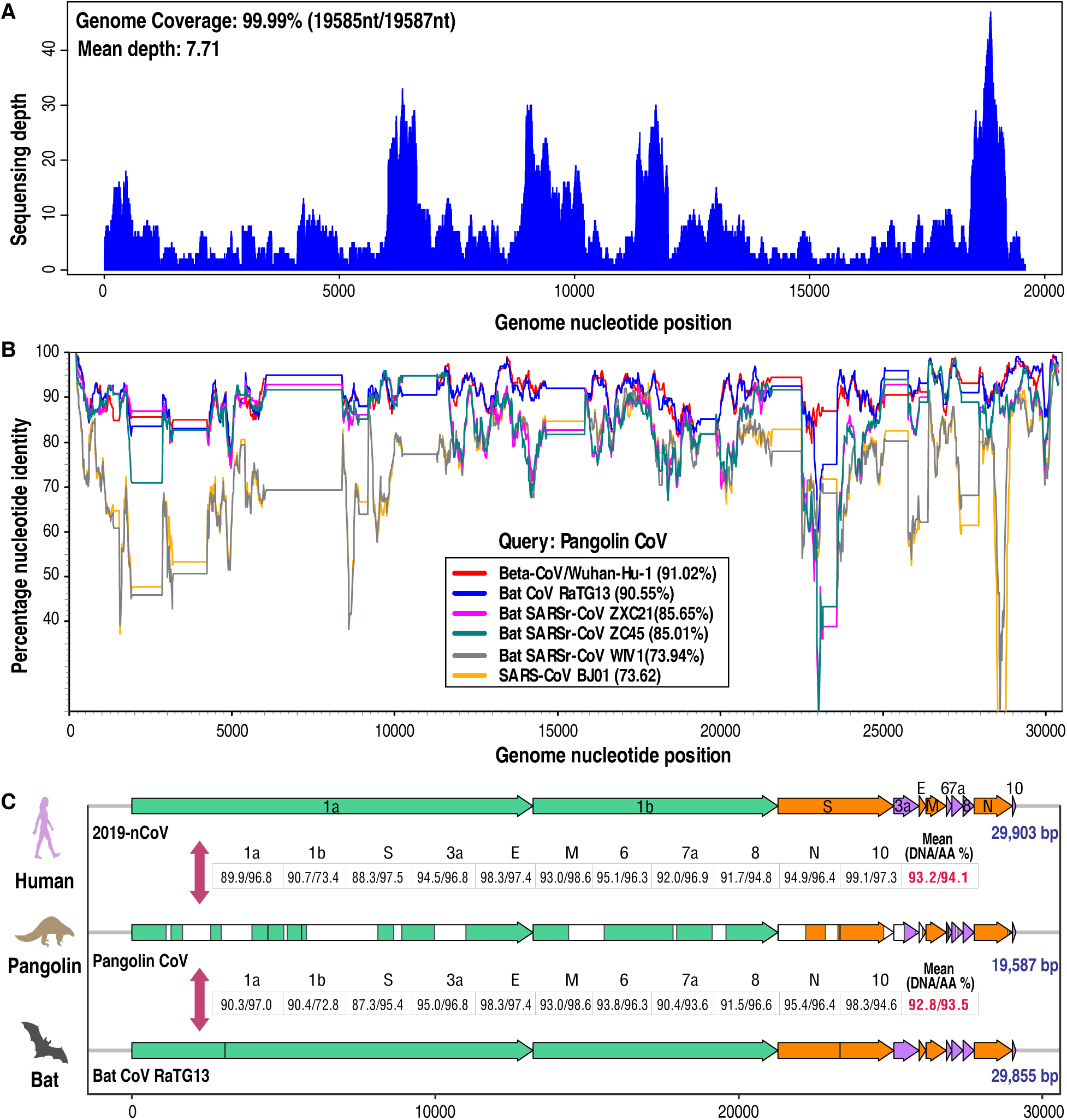
Genome-related analysis. (**A**) Sequence depth of mapped reads remapping to Pangolin-Cov. (**B**) Similarity plot based on the full-length genome sequence of Pangolin-Cov. Full-length genome sequences of 2019-nCoV (Beta-CoV/Wuhan-Hu-1), BatCov-RaTG13, Bat SARSr-CoV 21, Bat SARSr-CoV45, Bat SARSr-CoV WIV1, and SARS-CoV BJ01 were used as reference sequences. (**C**) Comparison of common genome organization similarity among 2019-nCoV, Pangolin-Cov and BatCov-RaTG13 linked to **Table S2**.

The viral genome organization of Pangolin-Cov was characterized by sequence alignment against 2019-nCoV (GenBank Accession MN908947) and RaTG13. The Pangolin-Cov genome consists of six major open reading frames (ORFs) common to coronaviruses and other four accessory genes (**Figure 1C** and **Table S2**). Further analysis indicates that Pangolin-Cov genes covered 2019-nCoV genes with coverage ranging from 45.8% to 100% (average coverage 76.9%). Pangolin-Cov genes shared high average nucleotide and amino identity with both 2019-nCoV (MN908947) (93.2% nt/94.1% aa) and RaTG13 (92.8% nt/93.5% aa) (**Figure 1C and Table S2**). Surprisingly, some of Pangolin-Cov genes showed higher aa sequence identity to 2019-nCoV than RaTG13, including orf1b (73.4/72.8), S-protein (97.5/95.4), orf7a (96.9/93.6), and orf10 (97.3/94.6). High S-protein amino acid identity implies function similarity between Pangolin-Cov and 2019-nCoV.

To determine the evolutionary relationships among Pangolin-Cov, 2019-nCoV and previously identified coronaviruses, we estimated phylogenetic trees based on the nucleotide sequences of the whole genome sequence, RNA-dependent RNA polymerase gene (RdRp), non-structural protein genes ORF1a and 1b, and the main structural proteins encoded by the S and M genes. In all phylogenies, Pangolin-CoV, RaTG13 and 2019-nCoV were clustered into a well-supported group, here named as “SARS-CoV-2 group” (**Figure 2** and **Figures S1 to S2**). This group represents a novel Beta-coronaviruses group. Within this group, RaTG13 and 2019-nCoV was grouped together, and the Pangolin-CoV was their lowest common ancestor. However, whether the basal position of the SARS-CoV-2 group is SARSr-CoV ZXC21 and/or SARSr-CoV ZC45 or not is still in debate. Such debate also occurred in both Wu et al.(*3*) and Zhou et al. (*2*)studies. Possible explanation is due to a past history of recombination in Beta-CoV group(*3*). It is noteworthy that our discovered evolutionary relationships of coronaviruses shown by the whole genome, RdRp gene, and S-gene were highly consistent with that discovered by complete genome information in Zhou et al. study(*2*). It indicates our Pangolin-CoV draft genome has enough genomic information to trace true evolutionary position of Pangolin-CoV in coronaviruses.

**Figure 2.**
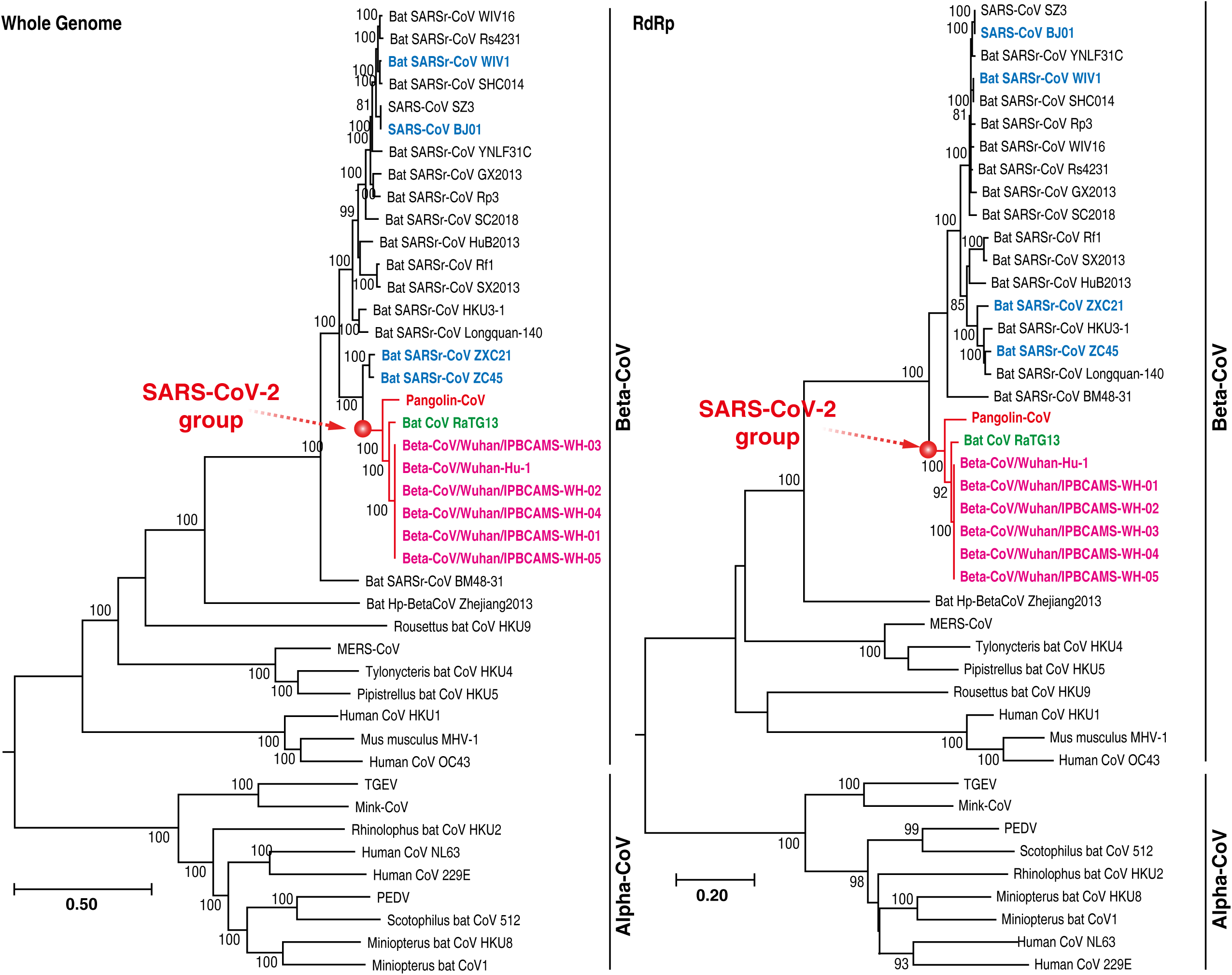
Phylogenetic relationship of coronavirus based on whole genome and RdRp gene nucleotide sequences. Red text denotes the Malayan Pangolin-CoV. Pink text denotes 2019-nCoV (SARS-CoV-2). Green text denotes a bat coronavirus having 96% similarity at genome level to SARS-CoV-2. Blue text denotes reference coronaviruses used in Figure1B. Detailed information can be found in **Materials and Methods**.

The coronavirus spike (S) protein consisting of 2 subunits (S1 and S2) mediates infection of receptor-expressing host cells and is a critical target for antiviral neutralizing antibodies. S1 contains a receptor binding domain (RBD) about 193 amino acid fragment, which is responsible for recognizing and binding with the cell surface receptor(*8, 9*). Zhou et al. had experimentally confirmed that 2019-nCoV is able to use human, Chinese horseshoe bats, civet, and pig ACE2 as an entry receptor in the ACE2-expressing cells(*2*), suggesting the RBD of 2019-nCoV mediates infection to human and other animals. To gain a sequence-level insight into understanding pathogen potential of Pangolin-Cov, we investigated the amino acid variation pattern of S1 protein from Pangolin-CoV, 2019-nCoV, RaTG13, and other representative SARS-CoVs. The amino acid phylogenetic tree showed the S1 protein of Pangolin-CoV is more closely related to that of 2019-CoV than RaTG13. Within the RBD, we further found Pangolin-CoV and 2019-nCOV was highly conserved with only one amino acid change (500H/500Q) (**Figure 3**) but not belonging to five key residues involved in the interaction with human ACE2(*2, 9*). In contrast, RaTG13 has 17 amino acid residue changes and 4 of them belonged to key amino acid residue (**Figure 3**). These results indicate Pangolin-Cov has similar pathogen potential to 2019-nCoV.

**Figure 3.**
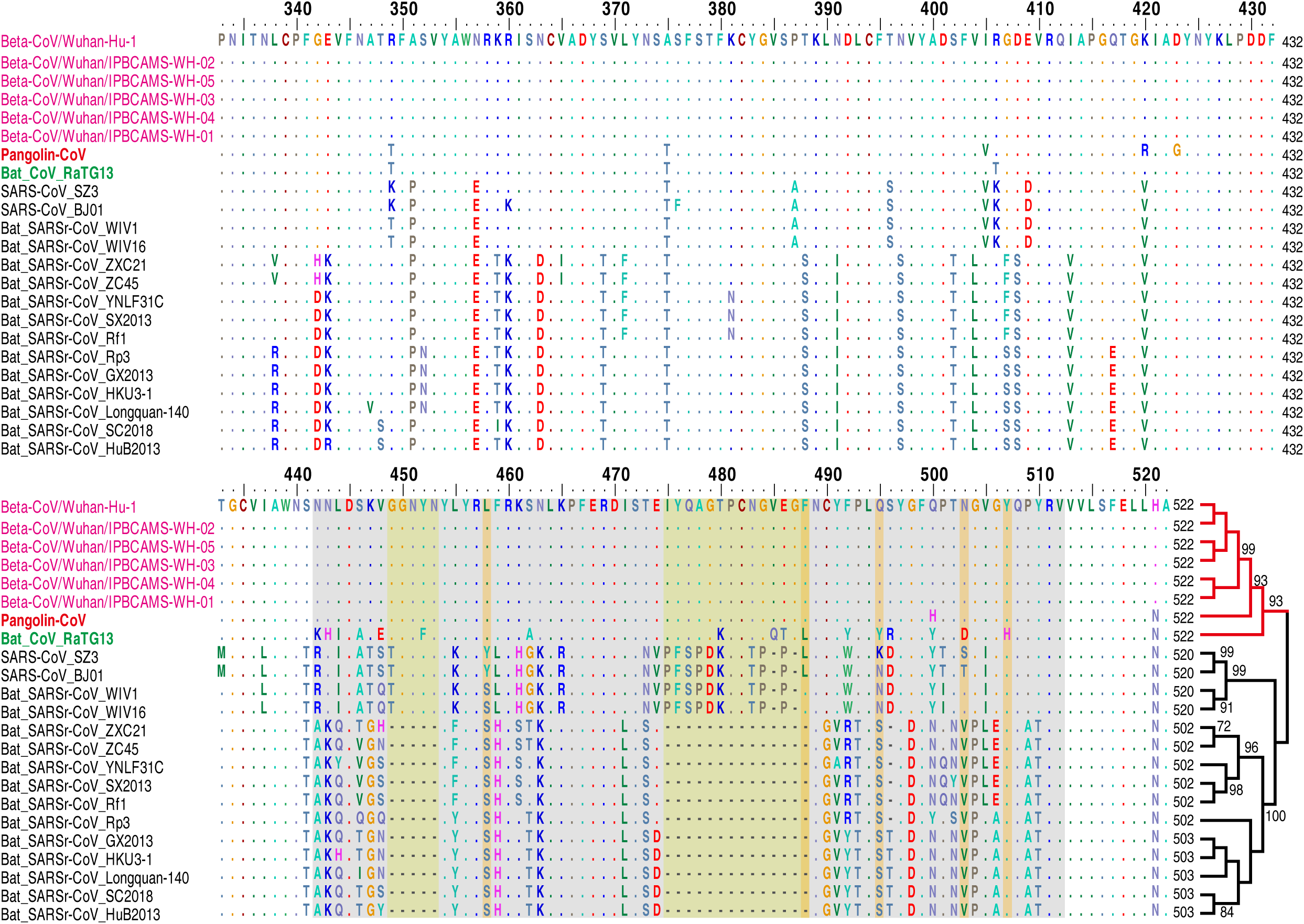
Amino acid sequence alignment of the S1 protein and its phylogeny. The receptor-binding motif of SARS-CoV and the homologous region of other coronaviruses are indicated by the grey box. The key amino acid residues involved in the interaction with human ACE2 are marked with the orange box. Bat SARS-like CoVs had been reported not to use ACE2, had amino acid deletions at two motifs marked by the yellow box. Detailed information can be found in **Materials and Methods**.

The nucleocapsid protein (N-protein) is the most abundant protein in coronavirus. The N-protein is a highly immunogenic phosphoprotein, and it is normally very conserved. The N protein of coronavirus is often used as a marker in diagnostic assays. To gain a further insight into the diagnostic potential for Pangolin-Cov, we investigated the amino acid variation pattern of N-protein from Pangolin-CoV, 2019-nCoV, RaTG13, and other representative SARS-CoV. Phylogenetic analysis based on N protein supports Pangolin-Cov as a sister taxon of 2019-nCoV and RaTG13 (**Figure 4**). We further found seven amino acid mutations can differentiate our defined “SAR-CoV-2 group” (12N, 26 G, 27S, 104D, 218A, 335T, 346N, 350Q) from other known SARS-CoVs (12S, 26D, 27N, 104E, 218T, 335H, 346Q, 350N). Two amino acid sites (38P and 268Q) are shared by Pangolin-Cov, RaTG13 and SARS-CoVs, which are mutated as 38S and 268A in 2019-nCoV. Only one amino acid residue shared by Pangolin-CoV and other SARS-CoVs (129E) is consistently changed in both 2019-nCoV and RaTG13 (129D). Our observed amino acid changes in N-protein would be useful for developing antigen for much more sensitive serological detection of 2019-nCoV.

**Figure 4.**
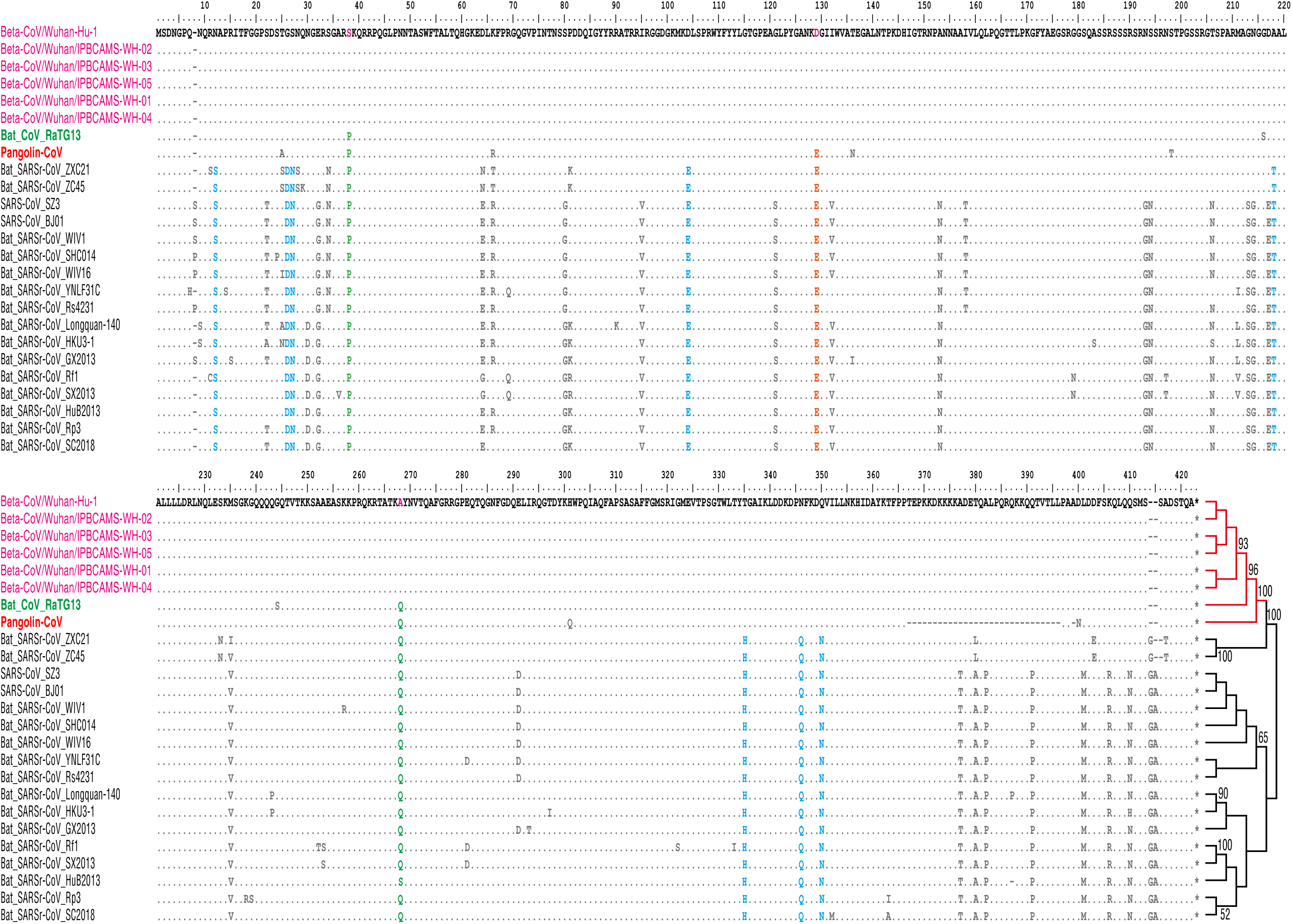
Amino acid sequence alignment of N protein and its phylogeny. Highly-conserved amino acid residues in N-protein marked by colors have diagnostic potential. Detailed information can be found in **Materials and Methods.**

Based on published metagenomic data, this study provides the first report on a potential closely related kin (Pangolin-CoV) of 2019-nCoV, which was discovered from dead Malayan Pangolins after extensive rescue efforts. Aside from RaTG13, the Pangolin-CoV is the most closely related to 2019-nCoV. Due to original sample unavailable, we did not perform further experiments to confirm our findings, including PCR validation, serological detection, and even the isolation of virus particle etc. However, on 7 February, researchers from the South China Agricultural University in Guangzhou reported pangolin would be the potential candidate host of 2019-nCoV for isolating a virus 99% similar to 2019-nCoV in genome (Data unpublished). Our discovered Pangolin-CoV genome showed 91.02% nt identity with 2019-nCoV, implying Pangolin-CoV could be different from that unpublished. Whether pangolin species is a good candidate for 2019-nCoV still need to be further investigated. Considering the wide spread of SARSr-CoV in their natural reservoirs, our findings would be meaningful to find novel intermediate hosts of 2019-nCoV for blocking interspecies transmission.

## Materials and Methods

### Data preparation

We downloaded raw data of lung08 and lung07 published by Liu’s study(*7*) from NCBI sequence read archive (SRA) under Bio Project PRJNA573298. Raw reads were first adaptor- and quality-trimmed using the Trimmomatic program (version 0.39)(*10*). For removing host contamination, Bowtie2 (version 2.3.4.3) (*11*)was used to map clean reads to the host reference genome of *Manis javanica* (NCBI Project ID: PRJNA256023). Only unmapped reads were mapped to 2019-nCoV reference genome (GenBank Accession MN908947) for identifying virus reads.

### Read Assembly and construction of consensus sequence

Virus-targeted reads were assembled *de novo* using MEGAHIT (v1.1.3)(*12*). Read remapping to assembled contigs was performed by using Bowtie2(*11*). Mapping coverage and depth were produced using Samtools (version 1.9)(*13*). Contigs were taxonomically annotated using BLAST 2.9.0+ against 2845 Coronavirus reference genomes (Table S1). Bat-Cov-RaTG13 genome was downloaded from NGDC database (https://bigd.big.ac.cn/) (Accession no. GWHABKP00000000)(*2*). 2019-nCoV reference genome was downloaded from NCBI (Accession no. MN908947)(*3*). Other coronavirus genomes were downloaded ViPR database (https://www.viprbrc.org/brc/home.spg?decorator=corona) on 6 February 2020. We further used reference-guided strategy to construct draft genome based on those contigs taxonomically annotated to 2019-nCoVs, SARS-CoV, and Bat SARS-like CoV. Each contig was aligned against 2019-nCoV reference genome with MUSCLE software (version 3.8.31)(*14*). Aligned contigs were merged into consensus scaffold with BioEdit version 7.2.5 (http://www.mybiosoftware.com/bioedit-7-0-9-biological-sequence-alignment-editor.html) following manually quality checking. Small fragments with less than length 25bp were discarded, if these fragments were not covered by any large fragments. The potential open reading frames (ORFs) of finally obtained draft genome were annotated by the alignment to 2019-nCoV reference genome (Accession no. MN908947).

### Phylogenetic relationship analysis

Sequence alignment was carried out using MUSCLE software(*14*). Alignment accuracy was checked manually base-by-base. Gblocks(*15*) was used to process the gap in aligned sequence. Using MegaX (version 10.1.7)(*16*), we inferred all maximum likelihood (ML) phylogenetic trees under the best-fit DNA/anima acid substitute model with 1000 bootstrap replications. Phylogenetic analyses were performed using the nucleotide sequences of various CoV gene data sets: the whole genome and ORF1a, ORF1b, Membrane (M) gene, spike (S) and RNA-dependent RNA polymerase (RdRp) gene. The best model of M is GTR+G and all others are GTR+G+I. Two additional protein-based trees were constructed under WAG+G (S1 subunit of S protein) and JTT+G (N-protein), respectively. Branches with values< 70% bootstrap were hidden in all phylogenetic trees.

## Acknowledgments

This study was supported by the Second Tibetan Plateau Scientific Expedition and Research (STEP) program (no. 2019QZKK0503), the National Key Research and Development Program of China (no. 2018YFC2000500), the Key Research Program of the Chinese Academy of Sciences (no. KFZD-SW-219), and the Chinese National Natural Science Foundation (no. 31970571).

---

**Figure S1.**
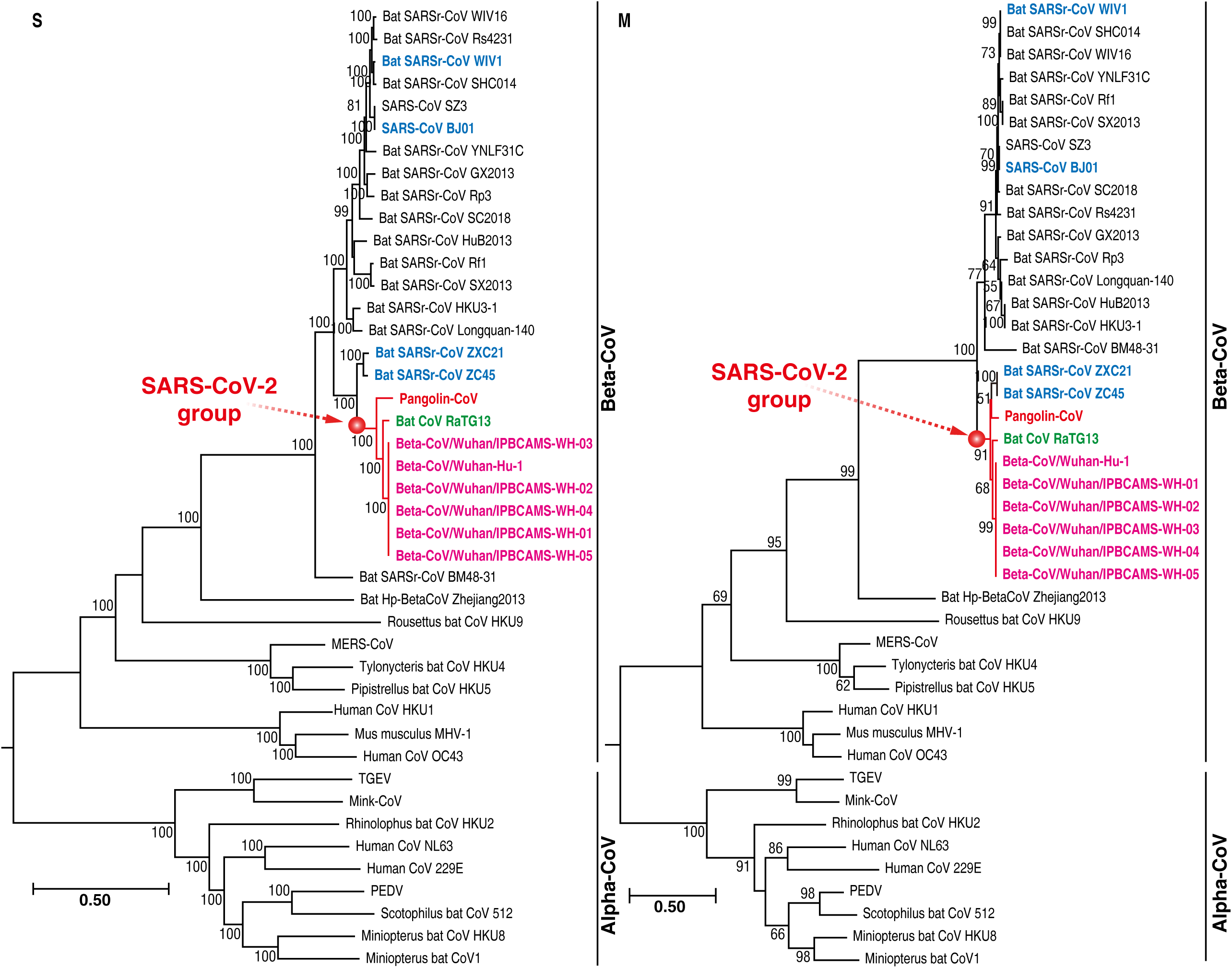
Phylogenetic relationship of coronavirus based on ORF1a gene (A) and ORF1b gene (B) nucleotide sequences.

**Figure S2.**
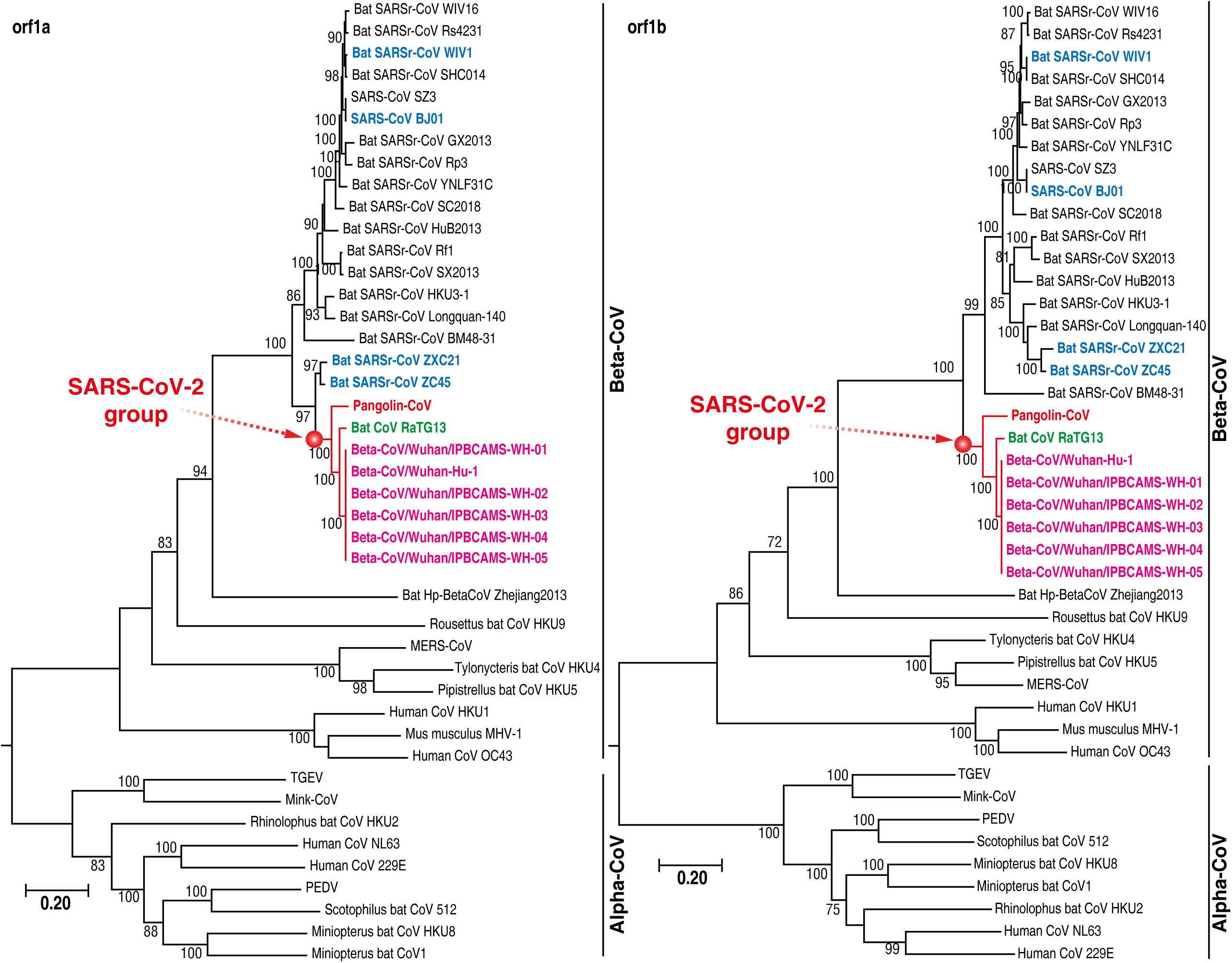
Phylogenetic relationship of coronavirus based on S gene (A) and M gene (B) nucleotide sequences.

**Table S1.**
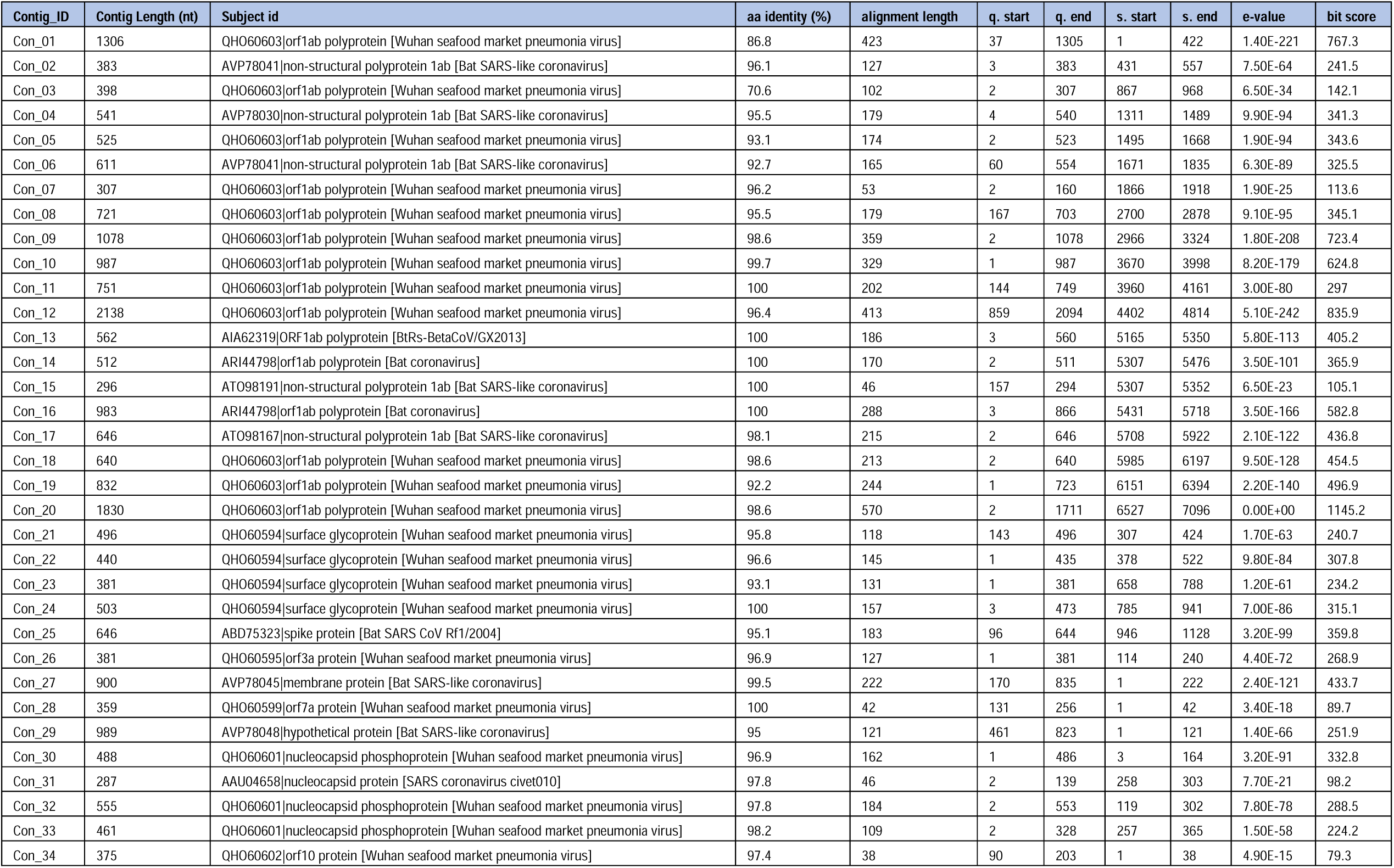
Contigs taxonomically annotated by using BLASTx against 2845 Coronavirus reference genomes.

**Table S2.** Comparing nt and aa sequence identity difference of ten genes among Pangolin-Cov, 2019-nCoV, and BatCov-RaTG13.

| Genes | 2019-nCoV (nt) | RaTG13 (nt) | Pangolin-CoV (nt) |
| --- | --- | --- | --- |
| orf1ab | 21290 | 21287 | 13669 (partial) |
| **orf1a** | 13128 | 13215 | 7342 (partial) |
| **orf1b** | 8072 | 8072 | 6327 (partial) |
| S | 3822 | 3810 | 2156 (partial) |
| 3a | 828 | 828 | 379 (partial) |
| E | 228 | 228 | 120 (partial) |
| M | 669 | 666 | 669 (Complete) |
| orf6 | 186 | 186 | 175 (partial) |
| orf7a | 366 | 366 | 296 (partial) |
| orf8 | 366 | 366 | 366 (Complete) |
| N | 1260 | 1260 | 1192 (partial) |
| orf10 | 117 | 117 | 117 (Complete) |
| **Nucleotide** | **2019-nCoV vs Pangolin-CoV** | **2019-nCoV vs RaTG13** | **Pangolin-CoV vs RaTG13** |
| orf1ab | 90.3% | 96.6% | 90.3% |
| **orf1a** | 89.9% | 96.2% | 90.3% |
| **orf1b** | 90.7% | 97.0% | 90.4% |
| S | 88.3% | 91.5% | 87.3% |
| 3a | 94.5% | 98.6% | 95.0% |
| E | 98.3% | 100.0% | 98.3% |
| M | 93.0% | 95.8% | 93.0% |
| orf6 | 95.1% | 98.8% | 93.8% |
| orf7a | 92.0% | 95.1% | 90.4% |
| orf8 | 91.7% | 96.9% | 91.5% |
| N | 94.9% | 96.7% | 95.4% |
| orf10 | 99.1% | 99.1% | 98.3% |
| Average | 93.2% | 96.9% | 92.8% |
| **Amino Acid** | **2019-nCoV vs Pangolin-CoV** | **2019-nCoV vs RaTG13** | **Pangolin-CoV vs RaTG13** |
| orf1ab | 87.1% | 95.8% | 86.9% |
| **orf1a** | 96.8% | 98.6% | 97.0% |
| **orf1b** | 73.4% | 92.2% | 72.8% |
| S | 97.5% | 96.7% | 95.4% |
| 3a | 96.8% | 100.0% | 96.8% |
| E | 97.4% | 100.0% | 97.4% |
| M | 98.6% | 100.0% | 98.6% |
| orf6 | 96.3% | 100.0% | 96.3% |
| orf7a | 96.9% | 96.9% | 93.6% |
| orf8 | 94.8% | 94.8% | 96.6% |
| N | 96.4% | 99.0% | 96.4% |
| orf10 | 97.3% | 97.3% | 94.6% |
| Average | 94.1% | 97.6% | 93.5% |

